# Mutating *PINK1* gene by paired truncated sgRNA/Cas9-D10A in Cynomolgus Monkey

**DOI:** 10.1101/2020.10.21.348862

**Authors:** Zhenzhen Chen, Jianying Wang, Yu Kang, Qiaoyan Yang, Xueying Gu, Dalong Zhi, Li Yan, Bin Shen, Yuyu Niu

## Abstract

Mutations of *PINK1* cause early-onset Parkinson’s disease (PD) with selective neurodegeneration in humans. However,current *PINK1* knockout mouse and pig models are unable to recapitulate the typical neurodegenerative phenotypes observed in PD patients, indicating that it is essential to generate *PINK1* disease models in non-human primates (NHPs) that are highly close to humans, to investigate the unique function of PINK1 in primate brains. Paired single guide RNA (sgRNA)/Cas9-D10A nickases and truncated sgRNA/Cas9, both enabling the reduction of the off-target effect without apparently compromising the on-target editing, are two optimized strategies in applying the CRISPR/Cas9 system to establish disease animal models. Here, we combined the two strategies and injected Cas9-D10A mRNA and two truncated sgRNAs into one-cell-stage cynomolgus zygotes to target *PINK1* gene. We show precise and efficient gene editing of the target site in the three new born cynomolgus monkeys. The frame shift mutations of *PINK1* in mutant fibroblasts leaded to the reduction of the mRNA, However, western-blot and immunofluorescencestaining confirmed the PINK1 proteinlevels werecomparable to that in the wild-type fibroblasts. We further reprogramed mutant fibroblast into induced pluripotent stem cells (iPSCs) and they show similar ability of differentiation into dopamine (DA) neurons. Taken together, our results showed that coinjection of Cas9-D10A nickase mRNA and sgRNA into one-cell-stage cynomolgus embryos enable the generation of human disease models in NHPs and the target editing by paired truncated sgRNA/Cas9-D10A in *PINK1* gene exon 2 did not impact the protein expression.

## Introduction

*PTEN-induced kinase I* (*PINK1*) was originally identified as a gene whose expression was up-regulated by overexpression of the tumor suppressor PTEN in carcinoma cell lines(Unoki et al., 2001). Within normal cells, the protein is mainly located in the mitochondria to protect cells from stress-induced mitochondrial dysfunction(Youle et al., 2012,Morais et al., 2014).Loss of function of PINK1 is closely associated with early-onset of Parkinson’s disease (PD)(Reed et al., 2019,Pickrell et al., 2015), a neurodegenerative disease characterized by the dysfunction of dopaminergic (DA) neurons in the substantia nigra (SN)(Goldstein et al., 2019,Poewe et al., 2017). The pathology of PD caused by *PINK1* mutationsis unclear, largely due to that the *PINK1* knock-out mouse and pig models were unable to recapitulate the typical PD-associated symptoms observed in human patients(Nakamura et al., 2007,Dawson et al., 2010,Zhou et al., 2015). Thus, it is necessary to generate a *PINK1* mutant animal model in non-human primates (NHPs),which show high similarity to human, to uncover the unique function of *PINK1* in primate brains.

CRISPR/Cas9-mediated disease models of NHPs containing precise genome modifications offer great hope for biomedical research(Lasbleiz et al., 2019,Vermilyea et al., 2018,Chan, 2013). However, the off-target effect of Cas9 on monkey genomes and other species is still inevitable, which limits its’ widespread applications(Zhang et al., 2015). Recent studies have demonstrated that CRISPR-Cas9 off-target effects can be reduced by increasing Cas9’s specificity including the use of high-fidelity Cas9 variants(Kleinstiver et al., 2016,Schmid-Burgk et al., 2020), or by reducing the generation of DSBs on off-target sites including the use of truncated sgRNAs or paired sgRNA/Cas9-D10A nickases(Bin et al., 2014,F Ann et al., 2013,Gopalappa et al., 2018). However, some of these methods often achieve lower off-target effects at the expense of reducing on-target efficiency. The Cas9 double nicking approach mediated by Cas9-D10A nickase requires the cooperation of two Cas9 nicking enzymes and a pair of sgRNAs to achieve DSBs. Unlike wild-type (WT) Cas9 that generates DSBs, Cas9-D10A nickase cuts only one strand of the DNA and generates single-stranded nick that can be repaired by the high-fidelity base excision repair (BER) pathway(Dianov et al., 2013), thus minimizing the introduction of indels. Several independent studies have clearly shown that the frequencies of off-target edits generated by paired Cas9-D10A nickases were lower than that with WT nucleases(Gopalappa et al., 2018,Guilinger et al., 2014,Ran et al., 2013).Moreover, the paired Cas9-D10A nickases exhibited comparable or sometimes significantly higher on-target efficiency than the individual WT Cas9.(Gopalappa et al., 2018). Because of the advantages offered by Cas-D10A, it has been applied to modify genes in Lactobacillus casei(Song et al., 2017), chicken DF1 cells(Lee et al., 2017) and pigs(Zeyland et al., 2018), all with alow off-target rate or even without off-target effect. However, the gene editing capacity of Cas9-D10A in NHPs have not been reported, also did truncated sgRNA.

Here in our study, we obtained three *PINK1* mutant cynomolgus monkeys via co-injection of Cas9-D10A mRNA and two truncated sgRNAs targeting exon 2 into one-cell-stage cynomolgus embryos.These*PINK1* mutant cynomolgus monkeys showed 12.5~100% editing efficiency on on-targets sites with no indels detected on the off-target sites tested, indicating paired truncated sgRNA/Cas9-D10A nickases enable the efficient and precise genome modifications in monkeys. The *PINK1* mutations did not affect PINK1 protein expression in fibroblasts derived from mutant monkeys compared to WT fibroblasts. Moreover, the DNA alterations of *PINK1* exon 2 didn’t affect the capacity of iPSCs reprogrammed from the mutant fibroblasts to differentiate into DA neurons. All these results suggest that paired truncated sgRNA/Cas9-D10A nickases could be applied to generate highly efficient and precise genome alterations in monkeys and targeting on *PINK1*exon 2 alone is not sufficient to model PD in NHP.

## Results

### Paired Truncated sgRNA/Cas9-D10A nickases Induces Efficient Genomic Editing in *PINK1* of Cynomolgus Embryos

To test whether the paired truncated sgRNA/Cas9-D10A system works in monkey embryos, we designed two truncated sgRNAs (Referred as S-gRNA and AS-gRNA) to target *PINK1* exon2, which encodes the kinase domain that is essential for PINK function (Figure 1A), and injected the mixture of 20ng/mL Cas9-D10A mRNA and 25 ng/mL sgRNA each into one-cell-stage zygotes of cynomolgus monkey. A total of 22 embryos normally developed to 8 cells (8C) or blastocyst stages were collected and examined for the presence of site-specific genome modifications by PCR, T7 endonuclease I (T7E1) cleavage assay and Sanger sequencing. As shown in Figure1B, 10 out of 22 (45.5%) embryos carries smaller ampliconor cleavaged bands after T7E1 digestion(Figure 1B).The TA cloning followed by Sanger sequencing analysis further showed that among the 10 edited embryos, 8 embryos contained 7~40-bp deletions with the editing efficiency varied from 9.1%~100%, one harbored 10-bp insertion and one carried both 4-bp deletion and 5-bp insertion(Figure 1C). Except for 27-bp deletion in embryo #13, all the indels detected in the edited embryos leaded to frame shift mutation, in principle, enabling the elimination of the full-length PINK1 protein production. In addition, to explore whether the microinjection of truncated sgRNA/Cas9-D10A RNAs impacts the monkey embryonic development, we calculated the development rate of 2-cell, 4-cell, 8-cell, morula and blastocyst stage respectively based on 55 Cas9-D10A injected embryos and 22 counterparts. For all the developmental stages tested, the develop rate of embryos with truncated sgRNA/Cas9-D10A injection was comparable to that in control group, thus demonstrating that microinjection of paired truncated sgRNA/Cas9-D10A into monkey embryonic cells didn’t affect early embryonic development(Fig 1D and Supplemental Fig 1A)

**Figure 1.**
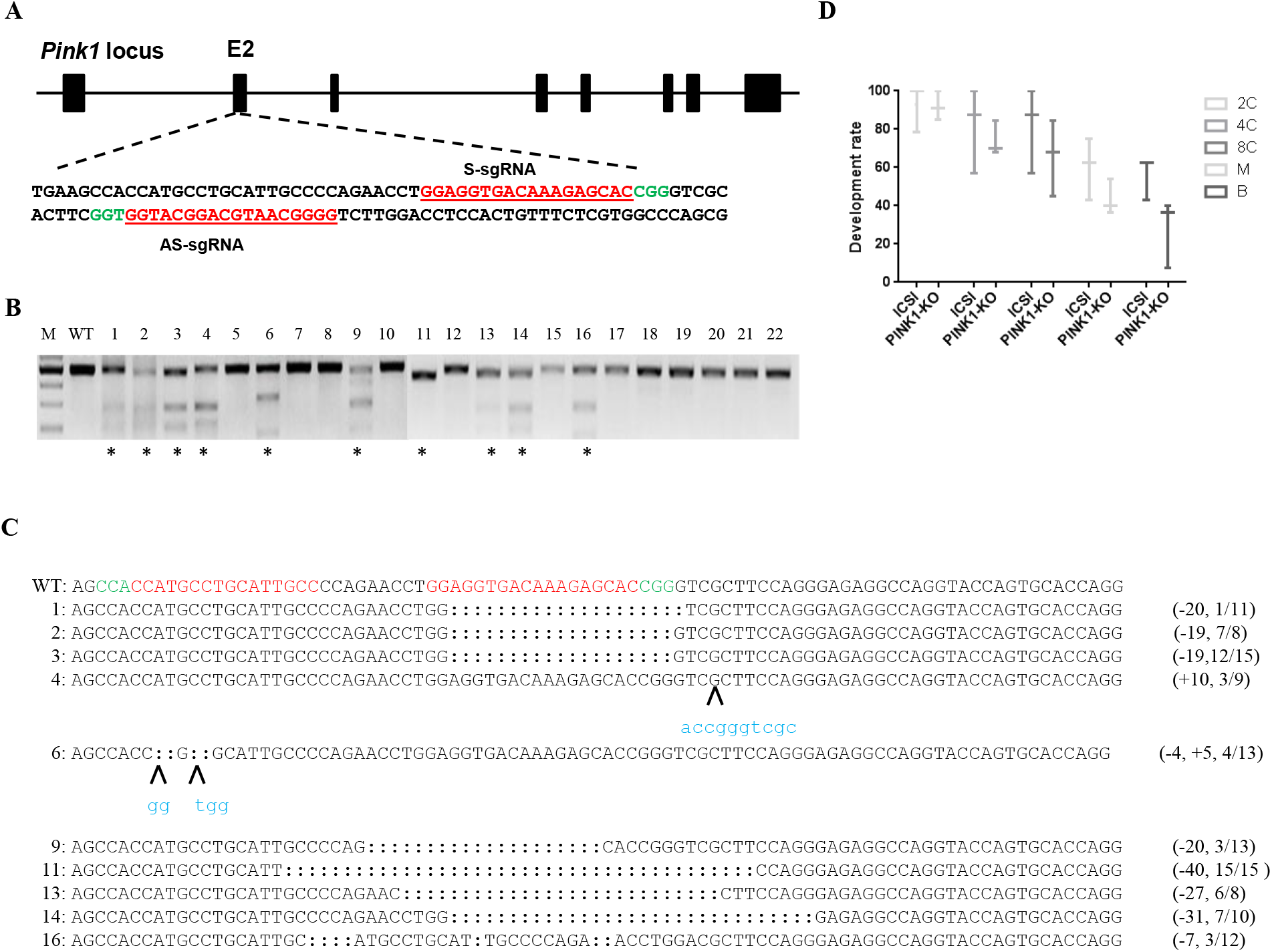
Paired Cas9-D10A nickases induce efficient genomeediting on *PINK1* in cynomolgus embryos. (A) Schematic diagram of sgRNAs targeting at PINK1 loci. Guide RNA sequences are underlined and highlighted in Red. PAM sequences are highlighted in Green. (B) Cas9-mediated on-target cleavage of PINK1 by T7E1 cleavage assay. PCR products were amplified and subjected to T7E1 digestion. Samples with cleavage bands were marked with an asterisk. (C) Editing profiles of marked samples in (B). Undigested PCR products from (B) were subjected to TA cloning. Single TA clones were picked and analyzed by DNA sequencing. For WT allele, the PAM sequences are highlighted in green and the sgRNA sequences are labeled in red; For the alleles with indels, the deleted bases are replaced with colons and the inserted bases are labelled with lower cases and highlighted in blue; deletions (-), and insertions (+). (D) Development rate in Cas9-D10A-injected embryos are comparable to that in ICSI-treated embryos.

### Paired truncated sgRNA/Cas9-D10A nickases Enable One-Step Generation of *PINK* Mutant Monkeys

Next, we set out to generate *PINK* modified cynomolgus monkeys using paired truncated sgRNA/Cas9-D10A. A total of 126 fertilized zygotes were microinjected with Cas9-D10A mRNA and sgRNA mixtures as described above. Total 51 Cas9-D10A-injected embryos that developed well were transferred into 25 surrogate females and six surrogates were successfully impregnated. Three of the six completed the pregnancy cycle (~150 days) and successfully gave birth to one infant or twins.(Figure 2A and Supplemental Figure 1B).By analysis of T7E1 assay on the DNA extracted respectively from ear or blood tissues, we found that three of the newborns carried *PINK1* mutations (referred as D10A-M1, -M2, and -M3, respectively), and one was unedited (referred as WT1)(Figure 2B). Next, to further dissect the on-target editing profiles and efficiency, we collected the DNA from fibroblasts derived from mutant monkeys, amplified the region around the target sites, performed TA cloning assay and sequenced 24 single clones for each newborn. The Sanger sequencing analysis showed that 12 out of 24 clones of 10A-M1 carried 8-bp deletion, and3 out of 24 clones of D10A-M2 had both 8-bp insertion and 19-bp deletion. For *PINK1* mutant monkey D10A-M3, half of the clones (12/24) harbored 19-bp deletion and the other half harbored 4-bp deletion, suggesting that D10A-M3 could be a homozygous mutant (Figure 2C).

The off-target effect is one of the major concerns for the CRISPR/Cas9 applications (Fu et al., 2013; Hsu et al., 2013; Pattanayak et al., 2013). To characterize the potential genome-wide off-target edits of truncated sgRNA/Cas9-D10A in newborn monkeys, a total of13off-targetsites (OTS)predicted by Cas-OFFinder were selected and the regions around all the potential off-target loci in mutant fibroblasts were PCR amplified followed by T7E1 assay and(or) DNA sequencing. No obvious indels was detected in all the potential off-target sites, suggesting that truncated sgRNA/Cas9-D10Awas able to decrease the productions of indels on the off-target sites in monkeys(Supplemental Figure 1C).

**Figure 2.**
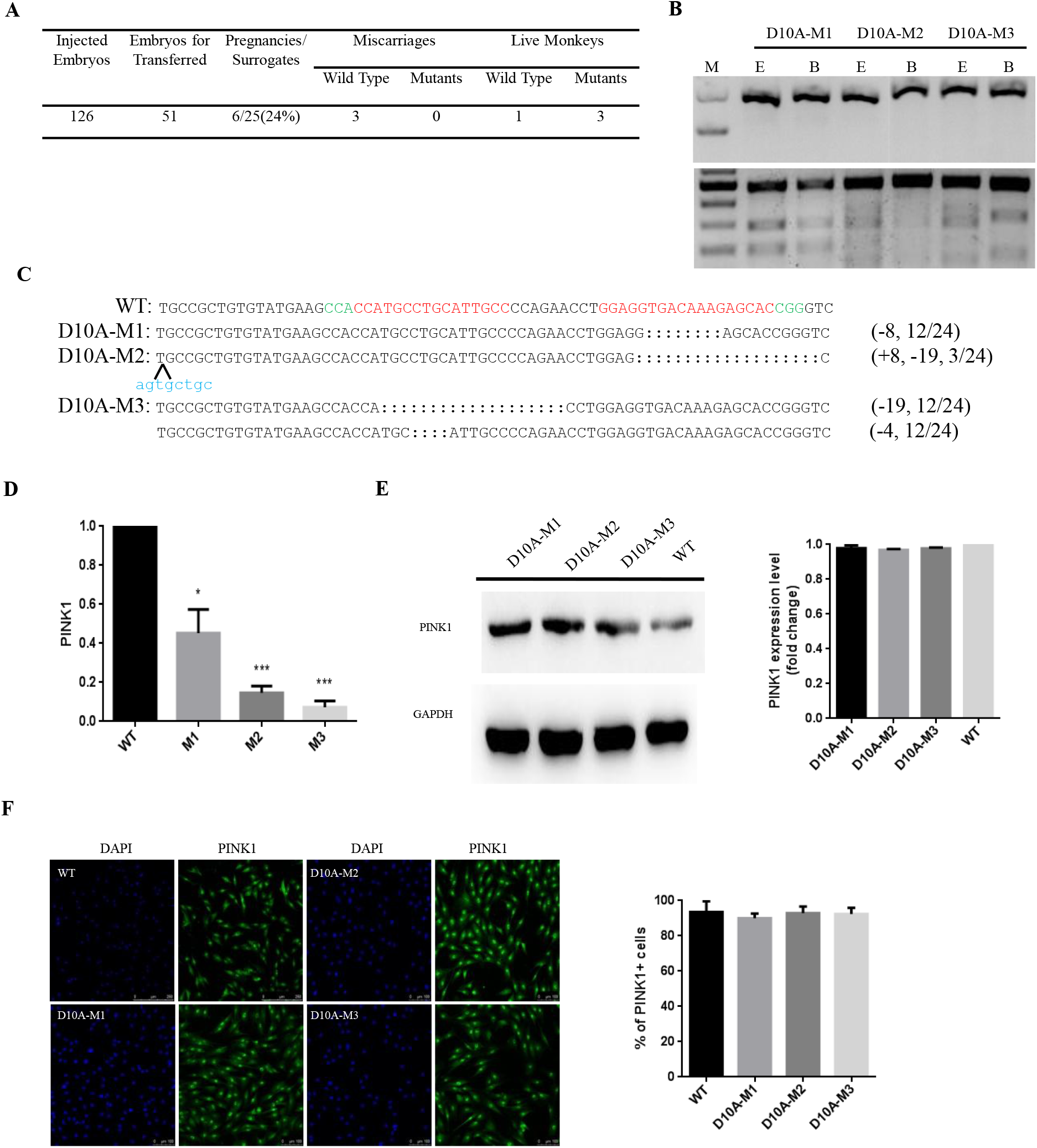
Paired Cas9-D10A nickases enable one-step generation of *PINK1* mutant monkeys. (A) Summary of embryos injected, transferred, impregnated and given birth. (B) T7E1 cleavage assay of target sites-containing DNA products amplified from ear or blood tissue of the mutant monkeys (D10A-M1, -M2 and -M3). The top panel represents the undigested PCR bands and the bottom panel stands for digested PCR products. E, ear; B, blood; M, marker. (C) Editing profiles of mutant monkeys. The region containing the target sites were amplified from mutant fibroblasts and the PCR products were subjected to TA cloning. Single TA clones were picked and analyzed by DNA sequencing. For WT allele, the PAM sequences are highlighted in green and the sgRNA sequences are labeled in red; For the alleles with indels, the deleted bases are replaced with colons and the inserted bases are labelled with lower cases and highlighted in blue; deletions (-), and insertions (+). (D) Western blotting assay on mutant and WT fibroblasts. The relative PINK1 expression levels were calculated via image J. Compared with WT monkey, all of the mutant monkeys exhibited similar PINK1 protein expression. (E) Representative images of Immunofluorescence staining on mutantand WT fibroblasts.The number of total cells and PINK1 positive cells were counted by image J.

To investigate whether the mutations of *PINK1* impact the expression levels of the messenger RNA (mRNA) and proteins, reverse transcription-PCR(RT-PCR) and Western blotting (WB) assays were respectively performed using the fibroblasts derived from *PINK1* mutant monkeys and counterparts. The *PINK1* mRNA levels in D10A-M1, -M2 and -M3 monkeys were respectively reduced to 45.3%, 14.7% and 7.3% of that in WT monkeys (Figure 2D), suggesting that the *PINK1* mutations on exon 2 probably induced the nonsense-mediated mRNA decay (NMD) via generating a premature stop codon. However, Western blotting analysis showed that the expression levels of PINK1 protein in mutant fibroblasts did not obviously reduced compared to that in WT fibroblasts (Figure 2E). The results of immunofluorescence staining also confirmed the presence of PINK1 residual protein in all the mutant fibroblasts (Fig 2F). All these results demonstrated that although all the *PINK1* mutant monkeys carried frame shift mutations on exon 2, the mutant fibroblasts didn’t show reduced expression of PINK1 protein compared to WT fibroblasts.

### The *PINK1* mutant fibroblasts-derived DA neurons does not show any specific PD-associated phenotypes

To further explore whether the *PINK1* mutations affect its’ function on DA neurons, we respectively reprogrammed the fibroblasts from D10A-M3 and WT monkeys into induced pluripotent stem cells (iPSCs) via Sendai virus-based method(Takahashi et al., 2006,Okita et al., 2007) and differentiated the selected iPS clones into DA neurons *in vitro*. A total of three iPS clones with *PINK1* mutations were selected and expanded for differentiation(Fig 3A–3B and Supplemental Fig 2A–2C). In order to identity the types of neuron cells differentiated from mutant and WT iPSCs, we stained the differentiated cells with specific markers of mature neurons (Tuj-1,NeuN, Foxa2 and Girk2),and DA neuron’s marker (TH). The result showed that the proportion of TH or Tuj-1 positive cells in mutant iPSC-DA neurons was comparable to that in WT iPSC-DA neurons (Fig 3C and supplemental Fig 2D–2E), indicating that the mutant *PINK1* did not influence the ability of iPSCs to differentiate into DA neurons.

**Figure 3.**
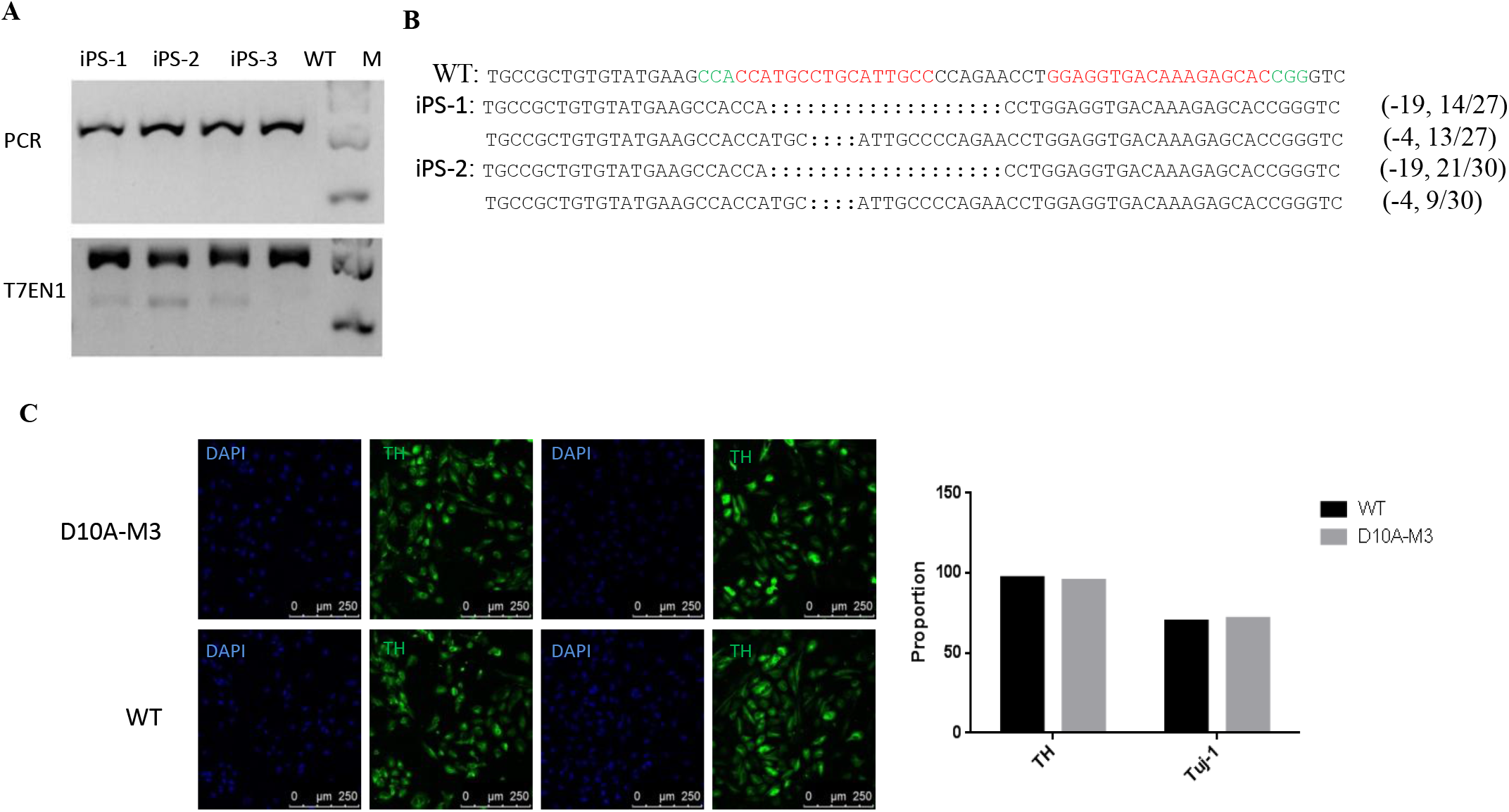
The *PINK1* mutant fibroblasts-derived DA neurons did not show any specific PD-associated phenotypes. (A) T7E1 cleavage assay of target sites amplified from three iPS clones reprogrammed from D10A-M3 fibroblasts. The top panel represents the undigested PCR bands and the bottom panel stands for digested PCR products. (B) Genotypes of iPS-1 and clone-2. (C) Representative images showing the DA neurons differentiated from mutant or WT iPSCs. Proportion of TH and Tuj-1 positive DA neurons from mutant and WT iPSCs were calculated. Scale bars, 250 um.

## Discussion

This study reported the first application of paired truncated sgRNA/Cas9-D10A nickases in modeling human diseases in NHPs. By injection of paired Cas9-D10A mRNA and two truncated sgRNAs targeting on *PINK1* exon 2 into one-cell-stage cynomolgus embryos, we successfully obtained three *PINK1* mutant cynomolgus monkeys with the editing efficiency varied from 9.1%~100%. Although the frame shift mutations were detected in mutant fibroblasts, the PINK1protein expression was not affected in contrast to that in WT fibroblasts due to a yet-unknown mechanism. Furthermore, the ability of iPSCs reprogrammed from fibroblasts to differentiate into PA neurons was not apparently impacted by the mutations of *PINK1* exon 2, suggesting *PINK1* exon 2 alone was probably not a good target for establishing *PINK1* knockout animal models.

Similar work was published earlier by Xiao-Jiang Li’s group(Yang et al., 2019), showing that CRISPR/Cas9 with paired sgRNAs enabled the generation of *PINK1* knockout rhesus models via inducing a large deletion across exon 2 and4. We speculate that the large deletions generated in Li’s group permitted the RNA splicing from *PINK1* exon 1 direct to exon 5, thus leading to non-sense mediated mRNA decay (NMD)(Brogna et al., 2009) and consequently eliminating protein translation. We also noticed that the *PINK1* mutant monkeys M3 and M4 generated by Xiao-Jiang Li’s group contained partial small frameshift deletion on either *PINK1* exon 2 or exon 4 and exhibited comparable protein levels compared to WT monkeys, a phenomenon consistent with our results. One potential possibility of this could be that the PINK WT protein stability or the capacity of *PINK* WT mRNA translation was enhanced upon the partial loss of PINK coding sequences. One the other hand, giving all the commercial antibodies used in our study were designed to recognize PINK1 carboxyl terminus, it cannot be ruled out that some yet-unidentified event, such as RNA mis-splicing, compensate the damage caused by the frame shift mutations on *PINK1* exon 2, thus eventually producing a protein that has a distinct N-terminus but an identical C-terminus compared to WT protein. Because of the presence of protein PINK1, all the mutant monkeys were born normally and are 6 years old now without any symptoms associated with PD. Because all the mutant monkeys are healthy so far, so we didn’t sacrifice them for further analysis.

Modeling human genetic diseases in NHPs via CRISPR/Cas9 technology hold great promise for studying human diseases and the development of corresponsive therapeutics, however, this process is challenged by the genome editing approaches giving the potential toxicity to embryos during microinjection of CRISPR/Cas9 components. We previously successfully generated gene-modified cynomolgus monkeys via co-injection of Cas9 mRNA and sgRNAs into one-cell-stage embryos (Niu et al., 2014), therefore firstly proving Cas9 is an efficient and reliable method for NHPs model generation. However, the off-target effect, one of the major concerns of utilizing CRISPR/Cas9 system for gene manipulation, is still unsolved. Double DNA nicking mediated by Cas9 nickases (D10A or H840A) and truncated sgRNAs for targeting are two optimal ways to minimize the generation of off-target edits without apparently impacting on-target editing efficiency(Ran et al., 2013,Fu et al., 2014). Here in our study, we proved that microinjection of Cas9 D10A mRNA and paired truncated sgRNAs into one-cell-stage cynomolgus embryos induced efficient gene modifications, enabled one-step generation of *PINK1* mutant monkeys and did not produce detectable indels on the top 13 potential off-target sites. All these results indicate that paired Cas9 could be an efficient and accurate approach to establish human diseases models in NHPs.

## Materials and Methods

### Ethics statement

Healthy cynomolgus monkeys (Macaca fascicularis), ranging in age from two to six years with body weights of four to six kg, were selected for use in this study. All animals were housed at the Yunnan Key Laboratory of Primate Biomedical Research (LPBR). All animal procedures were performed following the association for Assessment and Accreditation of Laboratory Animal Care International (AAALAC) for the ethical treatment of primates (approval number KBI K001115033-01).

### Preparation of mRNA and sgRNA

hCas9_D10A plasmids were obtained from Addgene (#41816). The plasmid was linearized with the restriction enzyme PmeI, and mRNA was synthesized and purified using an In Vitro RNA Transcription Kit (mMESSAGE mMACHINE T7 Ultra kit, Ambion). SgRNA oligos were amplified and transcribed in vitro using the GeneArt Precision gRNA Synthesis Kit (Thermo) and purified with the MEGAclear Kit (Thermo) according to the manufacturer’s instructions.

### Oocyte collection and in vitro fertilization

Oocyte collection and fertilization were performed as previously described(Niu et al., 2010). In brief, 10 healthy female cynomolgus monkeys aged 5–8 years with regular menstrual cycles were selected as oocyte donors for superovulation, which was performed by intramuscular injection with rhFSH (recombinant human follitropin alpha, GONAL-F, Merck Serono) for 8 days, then rhCG (recombinant human chorionic gonadotropin alpha, OVIDREL, Merck Serono) on day 9 Oocytes were collected by laparoscopic follicular aspiration 32–35 h after rhCG administration. Follicular contents were placed in Hepes-buffered Tyrode’s albumin lactate pyruvate (TALP) medium containing 0.3% BSA at 37 °C. Oocytes were stripped of cumulus cells by pipetting after a brief exposure (<1 min) to hyaluronidase (0.5 mg/mL) in TALP-Hepes to allow visual selection of nuclear maturity metaphase II (MII; first polar body present) oocytes. The maturity oocytes were subjected to intracytoplasmic sperm injection (ICSI) immediately and then cultured in CMRL-1066 containing 10% fetal bovine serum (FBS) at 37 °C in 5% CO2. Fertilization was confirmed by the presence of the second polar body and two pronuclei.

### sgRNA injection, embryo culture, and transplantation

Six to eight hours after ICSI, the zygotes were injected with a mixture of Cas9-D10A mRNA (20ng/mL) and sgRNA (25 ng/mL) with total volume 5 pL for each zygote. Microinjections were performed in the cytoplasm of oocytes using a microinjection system under standard conditions. Zygotes were then cultured in the chemically defined hamster embryo culture medium-9 (HECM-9) containing 10% fetal bovine serum (FBS, GIBCO) at 37 °C in 5% CaO_2_ to allow embryo development. The culture medium was replaced every other day until the blastocyst stage. The cleaved embryos with high quality at the two-cell to blastocyst stage were transferred into the oviduct of the matched recipient monkeys. Twenty five monkeys were used as surrogate recipients. The earliest pregnancy diagnosis was performed by ultrasonography about 20-30 days after the embryo transfer. Both clinical pregnancy and the number of fetuses were confirmed by fetal cardiac activity and presence of a yolk sac as detected by ultrasonography.

### Off-Target Analysis

All potential off-target sites with homology to the 23 bp sequence (sgRNA+PAM) were retrieved by a base-by-base scan of the whole Cynomolgus genome, allowing for ungapped alignments with up to four mismatches in the sgRNA target sequence. In the output of the scan, potential off-target sites with less than three mismatches in the seed sequence (1 to 7 base) were selected to PCR amplification using umbilical cord genomic DNA as templates. The PCR products were first subject to T7E1 cleavage assay. The potential off-target sites yielding typical cleavage bands were considered as candidates, then the PCR products of the candidates were cloned and sequenced to confirm the off-target effects.

### Western Blotting

For western blotting analysis, fibroblasts from ear was lysed, of which PINK1 protein expression were analyzed using GAPDH as an internal loading control for protein. About 40 mg of lysate was mixed with 5X loading buffer, boiled for 5 min and loaded onto 10% SDS-PAGE gels and transferred onto a PVDF membrane (Millipore). The membranes were blocked with 5% non-fat milk for 2 hr at room temperature, and then incubated with primary antibodies (PINK1, Cell Signaling Technology; GAPDH, Kangcheng) overnight at 4°C and subsequently for 1 hr at room temperature (25°C) with Goat Anti-Rabbit IgG Antibody, (H+L) HRP conjugate (Millipore) or Goat anti-Mouse IgG, (H+L) HRP conjugate (Millipore). The epitope was tested using Pierce ECL Western Blotting Substrate (Thermo Scientific).

### Reprogramming fibroblasts into iPSCs

Fibroblasts were infected by Sendai virus(Thermo Fisher Scientific, A16517) Transfected fibroblasts (approximately 1*105 cells per nucleofection) were directly plated into three 10cm feeder-seeded dishes in Dulbecco’s modified Eagle’s medium (DMEM) (Thermo Fisher Scientific, 11965-092), which contained 10% fetal bovine serum (FBS) (Hyclone, SH30396.03HI). The fibroblasts were replated 7 days post-infection and cultured in DMEM(Thermo Fisher Scientific,CT119955) with 10% FBS. The culture medium was changed every other day. On day 12 post-transfection, the medium was replaced with DMEM/F12 (Thermo Fisher Scientific, 10565-018) with15% KSR containing 10 ng/ml bFGF, 0.1mM β-Me, NEAA, 20% PS-Gro Medium(Stem RD). Colonies that morphologically resemble iPS colonies became gradually visible on day 15 after infection. iPS cell lines were cultured on X-ray-inactivated CF-1 mouse embryonic fibroblasts (MEFs) in PSC growth media.

### Dopaminergic neuron induction

iPS were digested with Collagenase IV (Gibco), and neural induction was induced by switching from ESC growth media to differentiation media in suspension culture [Advance DMEM/F12(Invitrogen): Neurobasal media(Invitrogen)(1:1 mixture) supplemented with 1 × N2 (Invitrogen), 1 × B27 (Invitrogen), 10 ng/ml bFGF (Millipore), 3 uM CHIR99021 (Cellagen Technology), 5 mMSB431542 (Cellagen Technology), 0.2 mMCompound E, and 0.1 mM LDN193189 (Cellagen Technology). After 6 days, EBs were transferred to 5 mg/ml laminin (Gibco)-coated plates for attachment culture, and the media were switched to NESCs culture media (Neurobasal media, including B27, N2, and NEAA (Sigma), 1% Glutmax (Sigma), 3 mM CHIR99021, 5 mM SB431542, 10 ng/ml bFGF, and 1,000 U/ml hLIF (Millipore)). To encourage cell propagation, 0.025% trypsin was used to digest NESCs when passaging. NESCs were routinely passaged to 1:8 to1:16 ratios every 3 to 4 days. On day 1, aspirate medium and replace with 2 mL of complete STEMdiff™ Dopaminergic Neuron Differentiation Medium. Incubate at 37°C and 5% CO2 for 5 - 6 days, performing a full medium change every other day with warm (37°C) complete STEMdiff™ Dopaminergic Neuron Differentiation Medium. On day 6/7, cells will reach 90 - 95% confluence.Seed cells onto pre-warmed (37°C) PLO/laminin-coated(0.05%, 5ug/ml, respectively) dish at a density of 4 x 10^4 - 6 x 10^4 cells/cm in complete STEMdiff™ Dopaminergic Neuron Differentiation Medium. Incubate at 37°C and 5% CO2 for 7 days, performing a full medium change every other day with warm (37°C) complete STEMdiff™ Dopaminergic Neuron Differentiation Medium.

On day 13/14, Seed dopaminergic neuronal precursors onto a pre-warmed (37°C) cell culture vessel coated with PLO/laminin at a density of 1.5 x 10^4 - 6 x 10^4 cells/cm2 in complete STEMdiff™ Dopaminergic Neuron Maturation Medium 1. Distribute cells evenly. Incubate at 37°C and 5% CO2, performing a full medium change every other day for 5 days. On day 18/19, replace medium with STEMdiff™ Dopaminergic Neuron Maturation Medium 2. Continue maturation of dopaminergic neurons in Medium 2 for a minimum of 2 weeks, performing a full medium change every other day.

### Immunostaining

The cells were fixed in 4% paraformaldehyde (DingGuo, AR-0211) at room temperature for 15 min and blocked with PBS (Corning, 21-040-CVR) that contained 0.2% Triton X-100 (Sigma-Aldrich, T8787) and 3% bovine serumAlbumin (BI, 04001A) at room temperature for 45 min. The cells were incubated with primary antibodies at 4 °C overnight. Secondary antibodies were incubated at room temperature for 1 hr. The nuclei were stained with DAPI (Roche Life Science, 10236276001). Antibody details were provided below.The following antibodies were used: anti-OCT4 (1:200; Abcam), Anti-Nanog (1:300; Abcam) and anti-SOX2 (1:200; Abcam).

## Acknowledgements

We acknowledge the members from animal facility of the Yunnan Key Laboratory of Primate Biomedical Research for excellent animal welfare and husbandry.

## Author contributions

Methodology: Z.Z. Chen, J.Y. Wang, Y. Kang, X.Y. Gu; Formal analysis: Z.Z. Chen; Data curation: Z.Z. Chen, Y. Kang, D.L. Zhi, L. Yan; Writing - review & editing: Z.Z. Chen, Y.Y. Niu, Q.Y. Yang, B. Shen; Visualization: Z.Z. Chen.; Supervision: Y.Y. Niu, B. Shen; Project administration: Y.Y. Niu; Funding acquisition: Y.Y. Niu(National Key Research and Development Program, 2016YFA0101401).

---

**Supplemental Fig 1.**
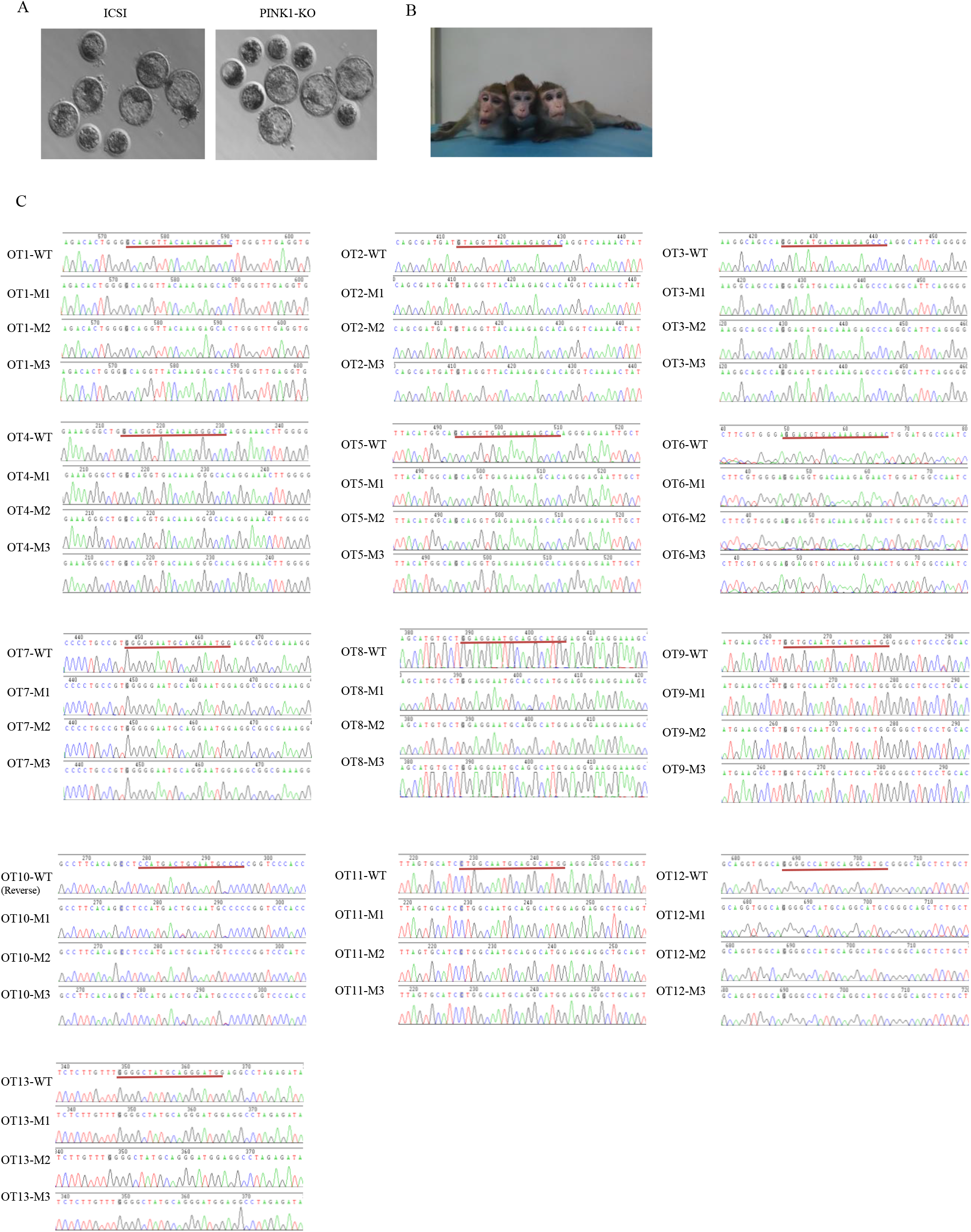
(A) Representative images of blastocysts developed from zygotes injected with or without Cas9-D10A mRNA and sgRNA. (B) Photos of three mutant monkeys. (C) Off-target sites analysis. The regions around the 13 potential OT sites were respectively amplified and subject to Sanger sequencing. Off-target sgRNAs were labelled by red horizontal lines and red arrowheads pointed to the mismatched bases.

**Supplemental Fig 2.**
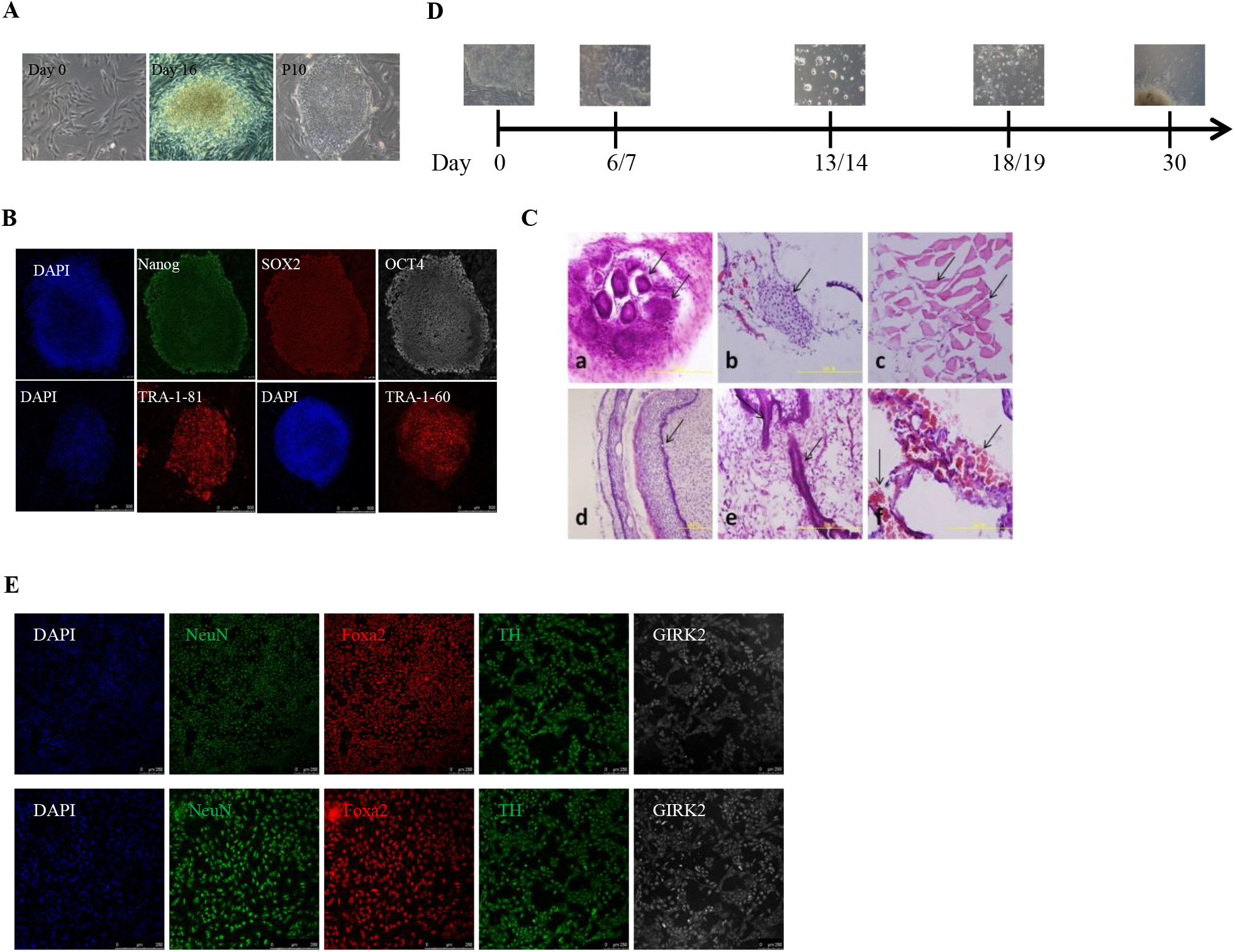
(A) Reprogramming fibroblasts into iPSCs by Sendai virus. The images represent the typic cell phenotypes observed at day 0 and day 16 after virus transduction and iPSC phenotypes at passage number 10. (B) Immunofluorescence staining of pluripotency marker—OCT4, Nanog, SOX2 and TRA-1-81/60 on selected iPS clones. Scale bars, 250 um. (C) Teratoma differentiation in immunodeficiency mice (NOD/SCID). a. Endoderm; b. Cartilage tissue; c. Muscle tissue; d. Phosphatid/columnar epithelium; e. Nervous tissue; f. Hematopoietic tissue. (D) Induction process from iPS to DA neurons.During the induction process, different differentiation and maturation media need to be replaced. About 30 days, DA neuron can be obtained. (E) Representative images showing immunofluorescent staining of DA neuron markers—NeuN, Foxa2, TH and GIRK2. Scale bars, 250 um.

